# An *Arabidopsis* AT-hook motif nuclear protein mediates somatic embryogenesis and coinciding genome duplication

**DOI:** 10.1101/2020.01.03.892869

**Authors:** Omid Karami, Arezoo Rahimi, Patrick Mak, Anneke Horstman, Kim Boutilier, Monique Compier, Bert van der Zaal, Remko Offringa

## Abstract

Plant somatic cells can be reprogrammed to totipotent embryonic cells that are able to form differentiated embryos in a process called somatic embryogenesis (SE), by hormone treatment or through overexpression of certain transcription factor genes, such as *BABY BOOM* (*BBM*). Here we show that overexpression of the *AT-HOOK MOTIF CONTAINING NUCLEAR LOCALIZED 15* (*AHL15*) gene induces formation of somatic embryos on *Arabidopsis thaliana* seedlings in the absence of hormone treatment. During zygotic embryogenesis, *AHL15* expression starts early in embryo development, and *AH15* and other *AHL* genes are required for proper embryo patterning and development beyond the heart stage. Moreover, *AHL15* and several of its homologs are upregulated and required for SE induction upon hormone treatment, and they are required for efficient *BBM*-induced SE as downstream targets of BBM. A significant number of plants derived from *AHL15* overexpression-induced somatic embryos are polyploid. Polyploidisation occurs by endomitosis specifically during the initiation of SE, assumingly due to AHL15-mediated heterochromatin decondensation coinciding with the acquisition of embryonic competency in somatic plant cells.

## Introduction

The conversion of somatic cells into embryonic stem cells is a process that occurs in nature in only a few plant species, for example on the leaf margins of *Bryophyllum calycinum* ^1^ or *Malaxis paludosa* ^2^, or from the unfertilized egg cell or ovule cells of apomictic plants ^3,4^. By contrast, for many more plant species, somatic cells can be converted into embryonic cells under specific laboratory conditions ^5,6^. The process of inducing embryonic cell fate in somatic plant tissues is referred to as somatic embryogenesis (SE). Apart from being a tool to study and understand early embryo development, SE is also an important tool in plant biotechnology, where it is used for asexual propagation of (hybrid) crops or for the regeneration of genetically modified plants during transformation ^7^.

SE is usually induced in *in vitro* cultured tissues by exogenous application of plant growth regulators. A synthetic analog of the plant hormone auxin, 2,4-dichlorophenoxyacetic acid (2,4-D), is the most commonly used plant growth regulator for the induction of SE ^8,9^. During the past two decades, several genes have been identified that can induce SE on cultured immature zygotic embryos or seedlings when overexpressed in the model plant *Arabidopsis thaliana* ^6,10^. Several of these genes, including *BABY BOOM* (*BBM*) and *LEAFY COTYLEDON 1* (*LEC1*) and *LEC2*, have now been recognized as key regulators of SE ^11–14^. Here we show that overexpression of *Arabidopsis AT-HOOK MOTIF CONTAINING NUCLEAR LOCALIZED 15* (*AHL15*) can also induce somatic embryos (SEs) on germinating seedlings in the absence of plant growth regulators. AT-hook motifs exist in a wide range of eukaryotic nuclear proteins, and are known to bind to the narrow minor groove of DNA at short AT-rich stretches ^15,16^. In mammals, AT-hook motif proteins are chromatin modification proteins that participate in a wide array of cellular processes, including DNA replication and repair, and gene transcription leading to cell growth, -differentiation, -transformation, - proliferation, and -death ^17^. The *Arabidopsis* genome encodes 29 AHL proteins that contain one or two AT-hook motifs and a PPC domain that directs nuclear localization and contributes to the physical interaction of AHL proteins with other nuclear proteins, such as transcription factors ^18,19^. *AHL* gene families are found in angiosperms and also in early diverging land plants such as *Physcomitrella patens* and *Selaginella moellendorffii* ^20,21^. *Arabidopsis* AHL proteins have roles in several aspects of plant growth and development, including flowering time, hypocotyl growth ^22,23^, flower development ^24^, vascular tissue differentiation ^25^, and gibberellin biosynthesis ^26^. How plant AHL proteins regulate these underlying biological events is largely unknown. Here we show that *AHL15* and its homologs play major roles in directing plant cell totipotency during both zygotic embryogenesis and 2,4-D- and BBM-mediated SE. Furthermore, our data show that AHL15 has a role in chromatin opening, and that its overexpression induces SE coinciding with endomitosis and polyploidy.

## Results

### Overexpression of *AHL* genes induces SE

To characterize the function of *AHL15*, we generated *Arabidopsis* lines overexpressing *AHL15* under control of the *35S* promoter (*35S:AHL15*). *AHL15* overexpression seedlings initially remained small and pale and then developed very slowly (Figure 1a). Three to four weeks after germination, seedlings from the majority of the transgenic lines (41 of 50 lines) recovered from this growth retardation (Figure 1a) and underwent relatively normal development, producing rosettes, flowers and finally seeds. However, in the remaining *35S:AHL15* lines (9 of 50 lines), globular structures could be observed on seedling cotyledons one- to two weeks after germination (Figure 1b). These structures developed into heart- or torpedo-shaped SEs (Figure 1c) that could be germinated to produce fertile plants.

**Figure 1.**
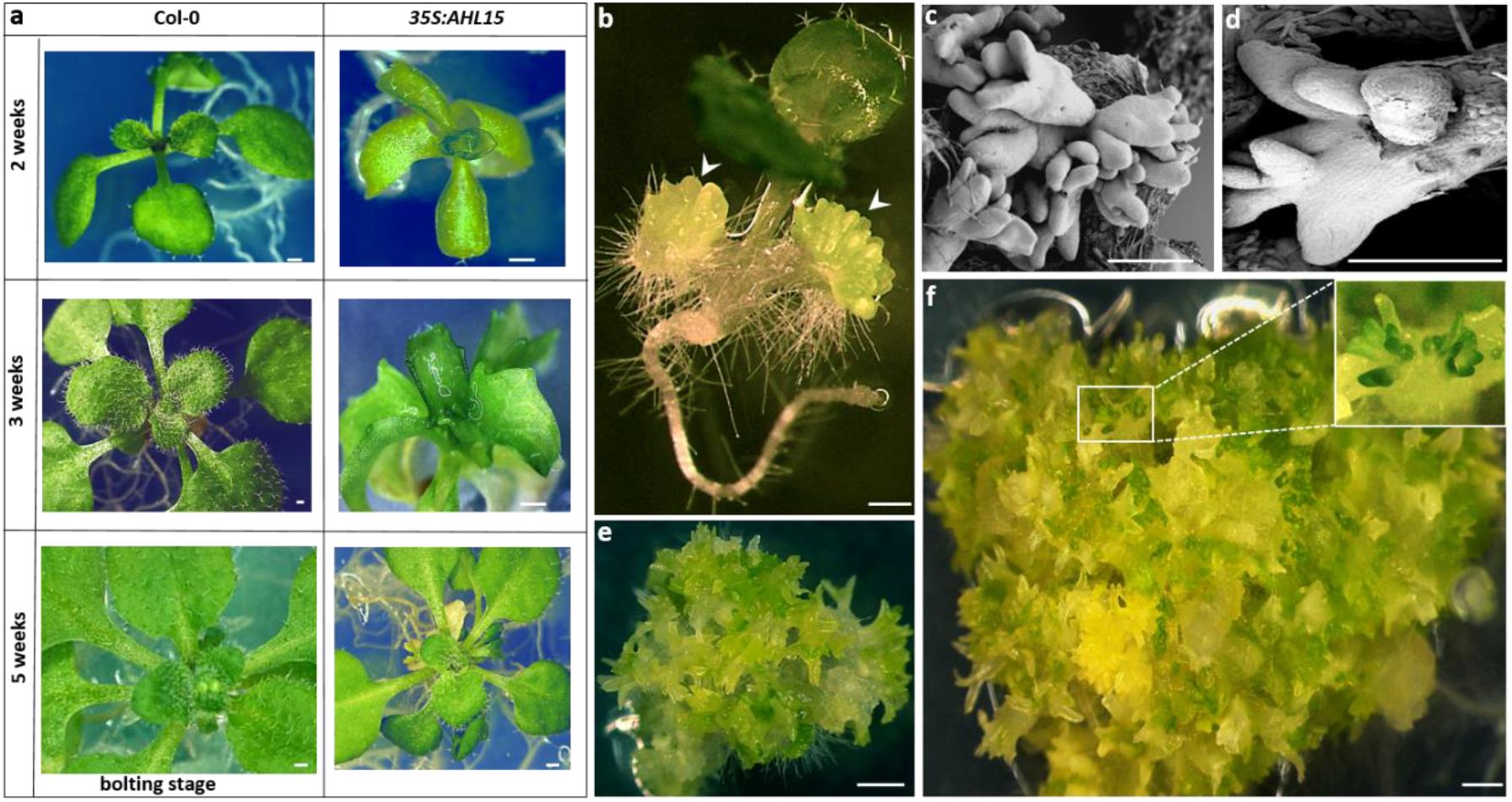
Overexpression of *AHL 15* delays *Arabidopsis* seedling development and induces SE. (a) The morphology of 2-, 3-, and 5-week-old wild-type and *35S:AHL15* plants grown in long day conditions (16 hr photoperiod). (b) Two-week-old *35S:AHL15 Arabidopsis* seedling with somatic embryos on the cotyledons (arrowheads). (c) Scanning electron micrograph showing torpedo stage somatic embryos on *35S:AHL15* cotyledons. (d) Scanning electron micrograph showing the secondary somatic embryos formed on a *35S:AHL15* primary somatic embryo. (e) The morphology of a 3-week-old overexpression *AHL15*-induced embryonic mass following secondary SE. (f) The morphology of a 2-month-old embryonic mass formed from a *35S:AHL15* seedling. Size bars indicate 0.5 cm in a and 1 mm in b-f.

In *Arabidopsis*, the cotyledons of immature zygotic embryos (IZEs) are the most competent tissues for SE in response to the synthetic auxin 2,4-dichlorophenoxyacetic acid (2,4-D) ^8^. Remarkably, a high percentage (85-95 %) of the IZEs from the selected *35S:AHL15* lines were able to produce somatic embryos when cultured on medium lacking 2,4-D. When left for a longer time on this medium, these primary *35S:AHL15* somatic embryos produced secondary somatic embryos (Figure 1d, e), and in about 20 % of the *35S:AHL15* lines, this repetitive induction of SE resulted in the formation of embryonic masses (Figure 1f). Overexpression of other *Arabidopsis AHL* genes encoding proteins with a single AT-hook motif (i.e. *AHL19*, *AHL20* and *AHL29*; Figure S1), did not induce SE on germinating seedlings, but did induce SE on a low percentage (20-30 %) of the IZEs in the absence of 2,4-D (Figure S2). These results suggest that AT-hook motif AHL proteins can enhance the embryonic competence of plant tissues, with AHL15 being able to most efficiently induce a totipotent state without addition of 2,4-D.

### *AHL* genes are important during zygotic embryogenesis

Given the role of *AHL* genes in promoting *in vitro* totipotency, we determined whether these genes also have a role during zygotic embryogenesis. Expression analysis using the *pAHL15:AHL15-GUS* and *pAHL15:AHL15-tagRFP* lines, both in the wild-type background, showed that *AHL15* is expressed in zygotic embryos (ZEs) from the 4-celled embryo stage onward, with its expression peaking at the bent-cotyledon stage (Figure 2a-i). In line with the previously reported nuclear localization of AHL proteins ^24,27^, the AHL15-tagRFP fusion protein was detected in the nucleus (Figure 2e-i and S3). Single and double *ahl15 ahl19* loss-of-function mutants carrying an artificial microRNA targeting *AHL20* (*ahl15 ahl19 amiRAHL20*) showed wild-type ZE development. Also *pAHL15:AHL15-GUS* plants produced wild-type embryos (Figure 2n) and seeds (Figure 2j). However, when we crossed these plants with the *ahl15* mutant, we were unable to obtain homozygous *ahl15 pAHL15:AHL15-GUS* seedlings among 50 F2 plants that we genotyped. Siliques of *ahl15/+ pAHL15:AHL15-GUS* plants contained brown, shrunken seeds (Figure 2k) that were unable to germinate. Moreover, around 25% of the embryos showed patterning defects leading to aberrantly shaped embryos that did not develop past the globular stage (Figure 2o). These results suggested that the AHL15-GUS fusion protein has a strong dominant negative effect on AHL function in the absence of the wild-type AHL15 protein. To confirm that the mutant phenotypes were caused by a dominant negative effect of the chimeric AHL15-GUS fusion protein, the *pAHL15:AHL15-GUS* construct was transformed to the *ahl15 pAHL15:AHL15* background. ZEs of *ahl15 pAHL15:AHL15* plants did not show any morphological defects (Figure 2l), and the resulting *ahl15 pAHL15:AHL15-GUS pAHL15:AHL15* siliques showed normal seed development (Figure 2m). These results indicate that wild-type AHL15 is able to complement the dominant negative effect of the AHL15-GUS fusion.

**Figure 2.**
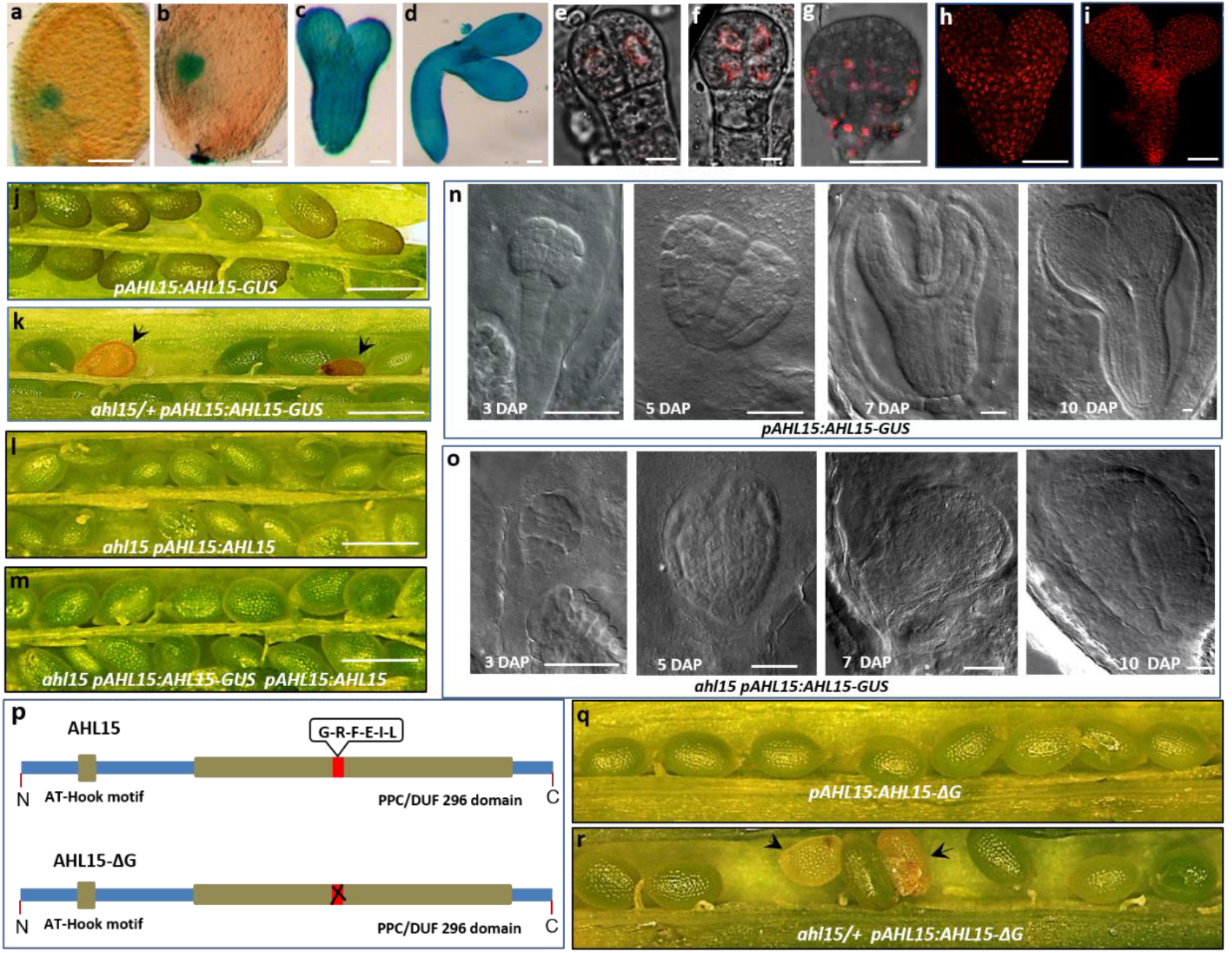
*AHL15* is expressed and essential during ZE. (a-d) Expression pattern of *pAHL15:AHL15-GUS* in globular- (a), heart- (b), torpedo- (c) and bent cotyledon (d) stage embryos. (e-i) Confocal microscopy images of *pAHL15:AHL15-tagRFP* four cell (e), 8 cell - (f), globular- (g), heart- (h), and torpedo- (i) stage embryos. (j) Wild-type seed development in a *pAHL15:AHL15-GUS* silique. (k) Aberrant seed development (arrowheads) in a *ahl15/+ pAHL15:AHL15-GUS* silique. (l,m) Wild-type seed development in *ahl15 pAHL15:AHL15* (l) or *ahl15 pAHL15:AHL15 pAHL15:AHL15-GUS* (m) siliques. (n, o) DIC images of zygotic embryo development in siliques of *pAHL15:AHL15-GUS* (n) or *ahl15/+ pAHL15:AHL15-GUS* (o) plants at 3, 5, 7 or 10 days after pollination (DAP). Size bar indicates 100 µm in a, b, b, g, h, l and 10 µm in E, F and 1mm j-m and 40 µm in n,o. (p) The schematic domain structure of AHL15 and the dominant negative AHL15-ΔG version, in which six-conserved amino-acids (Gly-Arg-Phe-Glu-Ile-Leu, red box) are deleted from the C-terminal PPC domain. (q) Wild-type seed development in *pAHL15:AHL15-ΔG* siliques. (r) Aberrant seed development (arrowheads) in ahl15/+ *pAHL15:AHL15-ΔG* siliques (observed in 3 independent *pAHL15:AHL15-ΔG* lines crossed with the *ahl15* mutant).

AHL proteins bind to each other through their PPC domain and form complexes with non-AHL transcription factors through a conserved six-amino-acid region in the PPC domain. In Arabidopsis, expression of an AHL protein without the conserved six-amino-acid region in the PPC domain leads to a dominant negative effect, as the truncated protein can still bind to other AHL proteins, but creates a non-functional complex ^19^. Based on this finding and a similar dominant-negative approach reported for human AT-Hook proteins ^28^, we deleted these six amino acids from the PPC domain of AHL15 (Figure 2p) and expressed it under the *AHL15* promoter (*pAHL15:AHL15-ΔG*) in wild-type plants. All plant lines generated with a *pAHL15:AHL15-ΔG* construct (n=20) were fertile and developed wild-type ZEs (Figure 2q). As for the *ahl15/+ pAHL15:AHL15-*GUS lines, we observed defective seeds in siliques in *ahl15/+ pAHL15:AHL15-ΔG* plants (Figure 2r) and were unable to obtain homozygous *ahl15 pAHL15:AHL15-ΔG* progeny, indicating that this genetic combination is also embryo lethal. These results provide additional support for the dominant negative effect caused by the AHL15-GUS fusion protein.

### *AHL* genes are required for 2,4-D- and BBM-induced SE induction

Next we investigated the contribution of *AHL* genes to 2,4-D-induced SE, by culturing IZEs from *ahl* loss-of-function mutants on medium containing 2,4-D. Only a slight reduction in SE induction efficiency was observed in the single *ahl15* loss-of-function mutant (Figure 3a), which stimulated us to examine the contribution of other *AHL* genes in this process. RT-qPCR analysis showed that *AHL15, AHL19* and *AHL20* expression was significantly upregulated in IZEs following seven days of 2,4-D treatment (Figure 3b). Moreover, analysis of *pAHL15*:*AHL15-GUS* expression in IZEs showed that *AHL15* expression was specifically enhanced in the cotyledon regions where somatic embryos are initiated (Figure 3c, d). IZEs from triple *ahl15 ahl19 amiRAHL20* mutants produced significantly less somatic embryos (Figure 3a) and also led to a relative increase in the number of abnormal somatic embryos (Figure 3e). Whereas *pAHL15:AHL15-GUS* IZEs showed a wild-type capacity to form somatic embryos in the presence of 2,4-D, the *ahl15* mutant in the *pAHL15:AHL15-GUS* reporter background resulted in a strong decrease in the embryogenic capacity of the IZE explants (Figure 3a). The majority of the IZEs were converted into non-embryogenic calli that were not observed in the other genotypes (Figure 3e). These results indicate that *AHL15* and several homologs are required for 2,4-D-induced somatic embryo formation starting from IZEs.

**Figure 3.**
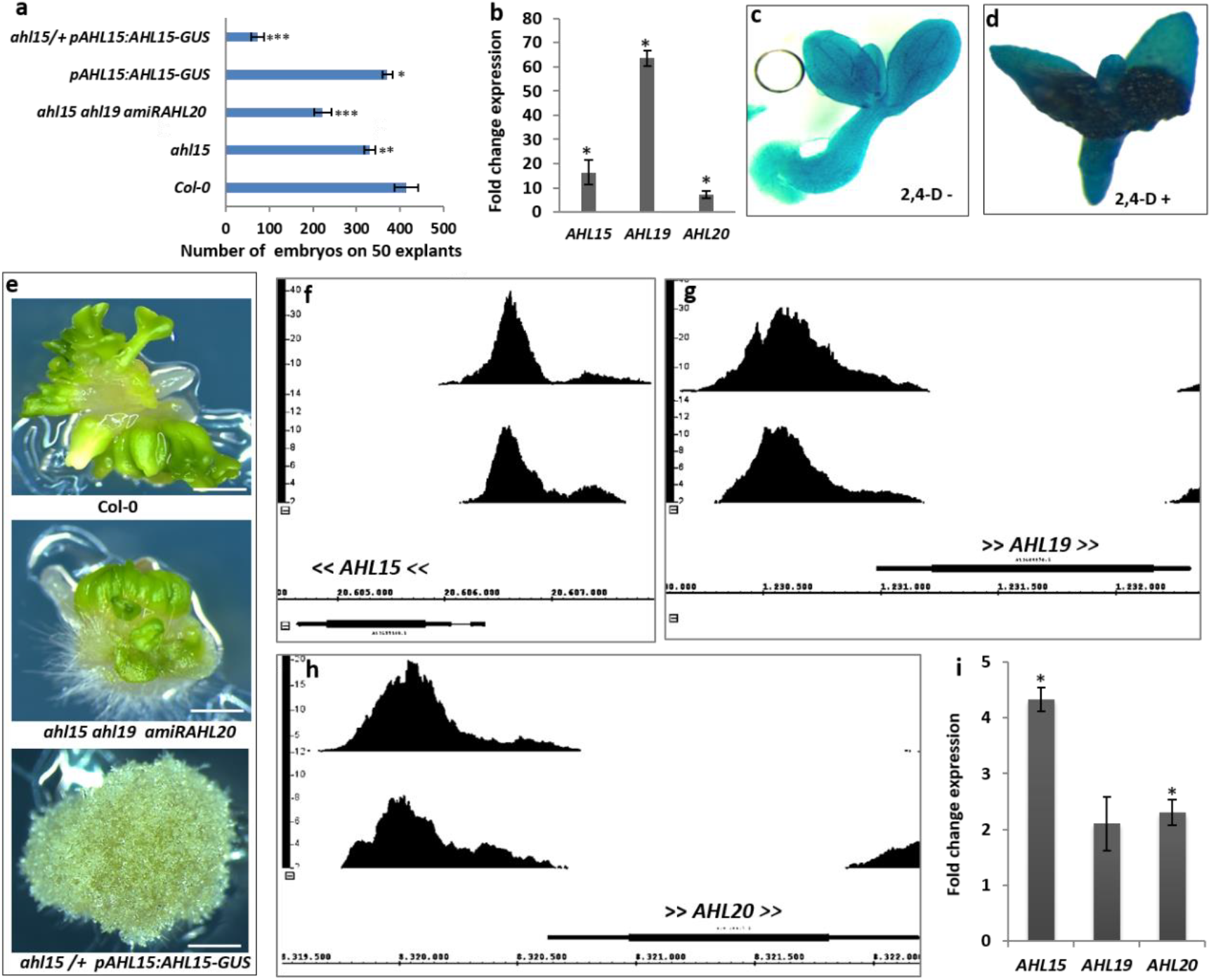
*AHL* genes are essential for 2,4-D- and BBM-induced SE. (a) The effect of *ahl* loss-of-function on the capacity to induce somatic embryos on IZEs by 2,4–D. Asterisks indicate statistically significant differences between the wild type and *ahl* mutant lines, as determined by the Student’s *t*-test (* p<0.05, ** p<0.01, *** p<0.001). Error bars indicate standard error of the mean of three biological replicates, with 50 IZEs per replicate. (b) RT-qPCR analysis of the fold change in expression of *AHL15*, *AHL19* and *AHL20* in IZEs cultured for 7 days on medium with 5 μM 2,4-D compared to medium without 2,4-D. Error bars indicate the standard error of the mean of four biological replicates. Asterisks indicate a significant enhancement of *AHL* gene expression in IZEs by 2,4-D treament (Student’s *t*-test, p < 0.001). (c, d) Expression of *pAHL15:AHL15-GUS* in IZEs cultured for 8 days in the absence (c) or presence (d) of 5 μM 2,4-D. (e) Embryo structures predominantly induced on IZEs obtained from wild-type (up), *ahl15 ahl19 amiRAHL20* (middle), or *ahl15/+ pAHL15:AHL15-GUS* (down) plants cultured for 2 weeks on 2,4-D medium. The genotype of *ahl15/+ pAHL15:AHL15-GUS* calli was verified by PCR analysis. Size bar indicates 1 mm. (f-h) ChIP-seq BBM binding profiles for *AHL15* (f), *AHL19* (g) and *AHL20* (h). The binding profiles from the *35S:BBM-GFP* (upper profile) and *pBBM:BBM-YFP* (lower profile) ChIP-seq experiments are shown. The x-axis shows the nucleotide position of DNA binding in the selected genes (TAIR 10 annotation), the y-axis shows the ChIP-seq score, and the arrow brackets around the gene name indicate the direction of gene transcription. (i) RT-qPCR analysis of the fold change in expression of *AHL* genes in DEX + CHX treated *35S:BBM-GR* seedlings relative to that in DEX + CHX treated Col-0 wild-type seedlings. Error bars indicate standard error of the mean of four biological replicates. Asterisks indicate a significant enhancement of *AHL* gene expression in DEX treated *35S:BBM-GR* seedlings compared to DEX treated Col-0 seedlings (Student’s *t*-test, p < 0.01).

Overexpression of the AINTEGUMENTA-LIKE (AIL) transcription factor BBM efficiently induces SE in in the absence of exogenous growth regulators ^13,29^. Genome-wide analysis of BBM binding sites using chromatin immunoprecipitation (ChIP) in 2,4-D and *35S:BBM*-induced somatic embryos showed that BBM binds to the promoter regions of *AHL15*, *AHL19* and *AHL20* (Figure 3f-h), suggesting that *AHL* genes are direct downstream BBM targets. To determine whether these genes are also transcriptionally regulated by BBM, we analyzed gene expression changes in *35S:BBM-GR* plants three hours after treatment with dexamethasone (DEX) and the translational inhibitor cycloheximide (CHX). These experiments showed that BBM activated the expression of *AHL15* and *AHL20*, but no statistically significantly difference in *AHL19* expression was observed (Figure 3i).

Next, we investigated the requirement for *AHL* genes in BBM-induced SE by transforming the *35S:BBM-GR* construct into the triple *ahl15 ahl19 amiRAHL20* mutant background. In wild-type Col-0, this construct induced SE in 40 of the 554 primary transformants (7 %), which was completely abolished in the *ahl15 ahl19 amiRAHL20* background (0 of the 351 primary transformants). These results, together with the observation that *AHL15* overexpression, like *BBM* overexpression induces spontaneous SE, suggest that induction of *AHL* gene expression is a key regulatory component of the BBM signaling pathway.

### AHL15 overexpression-mediated chromatin decondensation correlates with SE induction

Based on the observation in animal cells that AT-hook proteins are essential for the open chromatin in neural precursor cells ^30–32^, we investigated whether AHL15 modulates chromatin structure during SE initiation. Global chromatin structure is characterized by tightly condensed, transcriptionally-repressed regions, called heterochromatin, which can be visualized using fluorescent chromatin markers or DNA staining. Large-scale changes in heterochromatin in somatic plant cells are associated with cell identity reprogramming ^33,34^. Propidium iodide (PI) staining of chromosomal DNA in cotyledon cells of *35S:AHL15* IZEs showed a remarkable disruption of heterochromatin at between three and seven days after culture (Figure 4a). In contrast, cotyledon cells of wild-type IZEs did not show a clear change in heterochromatin state in this time period (Figure 4a). The *Arabidopsis* HISTONE 2B-GFP protein is incorporated into nucleosomes, providing a marker for the chromatin state in living cells ^34^. H2B-GFP fluorescence observations confirmed that the chromocenters in cotyledon cells of 7-day-cultured *35S::AHL15* IZEs (Figure 4b) were much more diffuse compared to cells of 3-day-cultured IZEs (Figure 4b). No significant differences in H2B-GFP signals were detected between cotyledon cells of three and seven day-incubated wild-type IZEs (Fig 4b). The number of detectable chromocentres was significantly decreased in cotyledon cells *35S:AHL15* IZEs cultured for 7 days relative to wild-type IZEs (Figure 4c). This result suggests that AHL15 promotes heterochromatin decondensation. Surprisingly, in cells expressing both *pAHL15:AHL15-tagRFP* and *pH2B:H2B-GFP* reporters, *AHL15-tagRFP* did not co-localize with the chromocenters (Figure S4), but showed a more diffuse nuclear distribution, suggesting that AHL15 regulates global chromatin decondensation.

**Figure 4.**
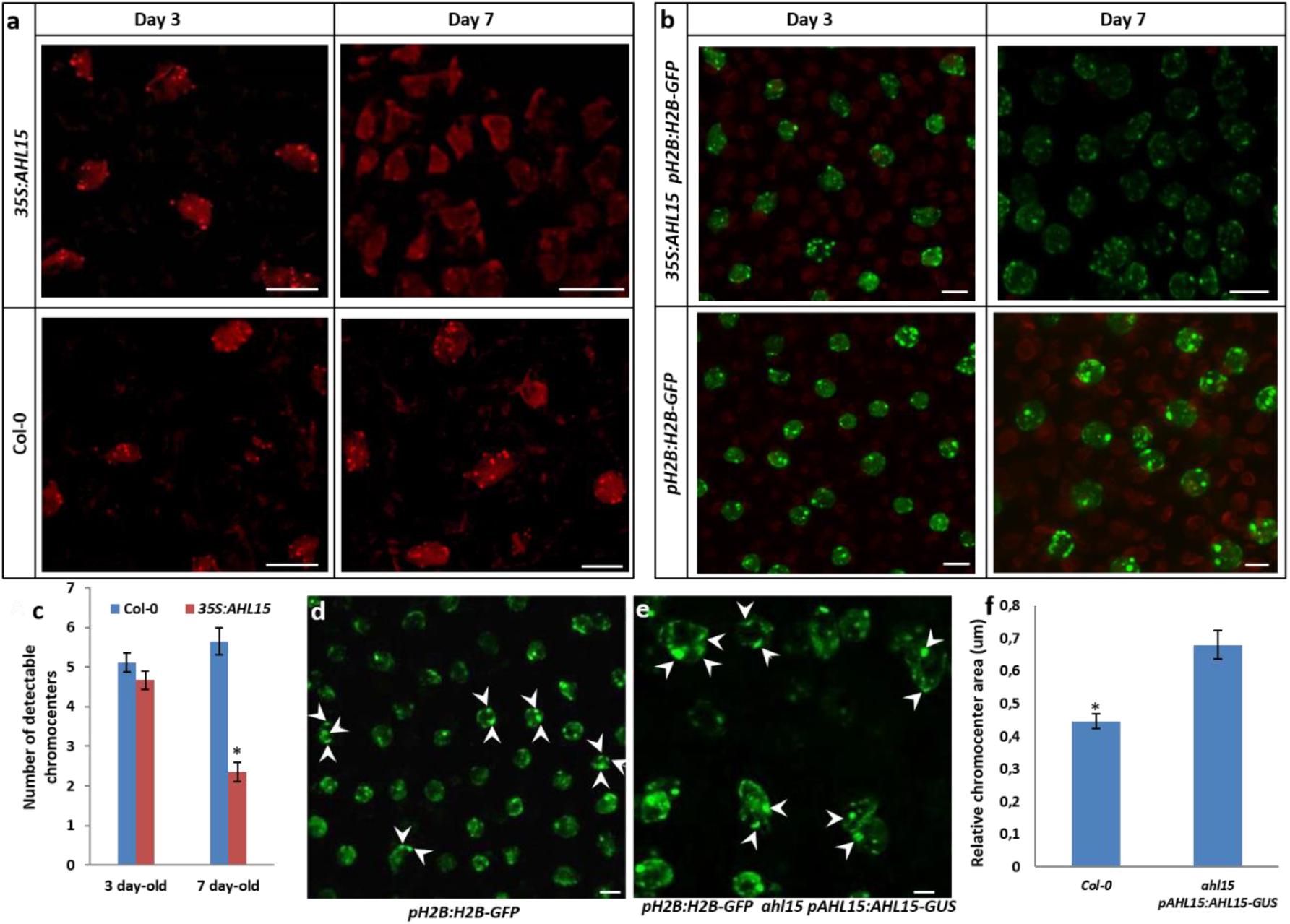
*AHL15* reduces heterochromatin condensation and chromocenter size. (a, b) Visualization of DNA compaction using propidium iodide (PI) staining (a) or H2B-GFP labelling (b) in cotyledon cell nuclei of wild-type and *35S::AHL15* IZEs 3 or 7 days after culture on B5 medium. Size bar indicates 6 µm in A and B. (c) Quantification of the number of conspicuous chromocenters labelled with PI in cotyledon cell nuclei of wild-type and *35S:AHL15* IZEs 3- or 7 days after culture on B5 medium. Error bars indicate standard error of the mean of 4 biological replicates. For each replicate 10 nuclei were analyzed. Asterisk indicate statistically significant differences (Student’s *t*-test, * p<0.01). (d, e) Visualization of chromocenters using the *pH2B:H2B-GFP* reporter in wild-type (d) and defective *ahl15 pAHL15:AHL15-GUS* ZEs (e) at 6 DAP. Size bar indicates 3.5 µm in d and e. (f) Quantification of the chromocenter area labelled with H2B-GFP in nuclei of wild-type and *ahl15 pAHL15:AHL15-GUS* ZEs at 6 DAP. Error bars indicate standard error of the mean of 6 biological replicates. For each replicate 30 chromocenters were measured (2 or 3 of the most clear chromocenters per nucleus), indicated with arrow heads in d and e). Asterisk indicates statistically significant differences (Student’s *t*-test, * p<0.01).

To explore whether the observed heterochromatin decondensation in *35S:AHL15* IZEs also occurs in other SE systems, we investigated the heterochromatin state in embryonic cells induced by 2,4-D. The H2B-GFP signals in cotyledon cells of *pH2B:H2B-GFP* IZEs cultured for 7 days on medium containing 2,4-D displayed moderate decondensation of heterochromatin compared to cells of 3-day-cultured IZEs, but did not show such diffuse heterochromatin (Figure S5a,b) as observed in *35S:AHL15* IZEs (Figure 4b). This suggests that the AHL15-induced chromatin decondensation is not a general trait of cells undergoing SE, but rather, is specific for AHL15-overexpressing cells. Altthough 2,4-D strongly upregulates *AHL15* expression during somatic embryogenesis (Figure 3b-d), *35S* promoter-driven *AHL15* expression induces to approximately 3-fold higher RNA levels (Figure S5c). This difference in expression levels might explain the strong chromatin decondensation observed in *35S::AHL15* IZEs compared to 2,4-D-treated IZEs.

To obtain insight into the role of AHL15 in chromatin decondensation during zygotic embryogenesis, we introduced the *pH2B:H2B-GFP* reporter into the *ahl15/+ pAHL15:AHL15-GUS* background. In defective *ahl15 pAHL15:AHL15-GUS* embryos, we observed irregular shaped chromocenters that were much larger than those in wild-type cells (Figure 4d-f). This result, together with the reduced heterochromatin condensation observed in cotyledon cells of *35S:AHL15* IZEs, suggests that *AHL15* plays a role in regulating the chromatin architecture during embryogenesis.

A previous study showed that blue light induces heterochromatin condensation in Arabidopsis cotyledons ^34^. We therefore analyzed the effect of blue light on *35S:AHL15*- and 2,4-D-induced SE. In both systems, incubating IZEs in blue light strongly inhibited SE (Figure S6a). Comparison of the heterochromatin state in IZE cotyledon cells using the H2B-GFP reporter revealed a significant increase of H2B-GFP signal in blue light-treated compared to white light-treated *35S: AHL15* cotyledons (Figure S6b). From this experiment we conclude that *AHL15* overexpression-mediated heterochromatin decondensation correlates with, and is probably required for its capacity to induce SE.

### *AHL15* overexpression induces polyploidy during SE initiation

Plants regenerated from somatic embryos derived from *35S*:*AHL15* IZEs (without 2,4-D treatment) developed large rosettes with dark green leaves and large flowers (Figure S7a). These phenotypes were not observed in *35S:AHL15* progeny obtained through zygotic embryogenesis. As these phenotypes are typical for polyploid plants, we determined the ploidy level of the plants. The number of chloroplasts in guard cells ^35^ of plants showing large flowers was two times higher (8-12) than that of diploid wild-type plants (4-6) (Figure S7b). Polyploidisation is also correlated with an increase in cell- and nuclear size in *Arabidopsis* and many other organisms ^36^. Indeed root cells of tetraploid *35S::AHL15* seedlings showed a larger nucleus and a larger cell volume than diploid control plants (Figure S7c), explaining the larger organ size observed for these plants. We used the centromere-specific HISTONE3-GFP fusion protein (*CENH3-GFP*) to count the number of chromosomes per cell ^37,38^. Seven to eight CENH3-GFP-marked centromeric dots could be detected in root cells of wild-type Arabidopsis plants and diploid *35S:AHL15* SE-derived plants (Figure S7d). By contrast, around 12-16 centromeric dots were observed in the larger nuclei in root cells of tetraploid *35S:AHL15* plants (Figure S7d). This confirmed that the plants with large organs that were regenerated from *AHL15* overexpression-induced somatic embryos are polyploid. Additional flow cytometry analysis on SE-derived plant lines confirmed that most of these plants were tetraploid, and two were octoploid (Table 1). The frequency of SE-derived polyploidy varied per *35S:AHL15* line, ranging from 18 to 69% (Table 1). No polyploid plants were obtained from somatic embryos induced by 2,4-D on wild-type IZEs, or by *BBM* overexpression in seedlings (Table 1), indicating that polyploidisation is specifically induced by *AHL15* overexpression.

**Table 1.**
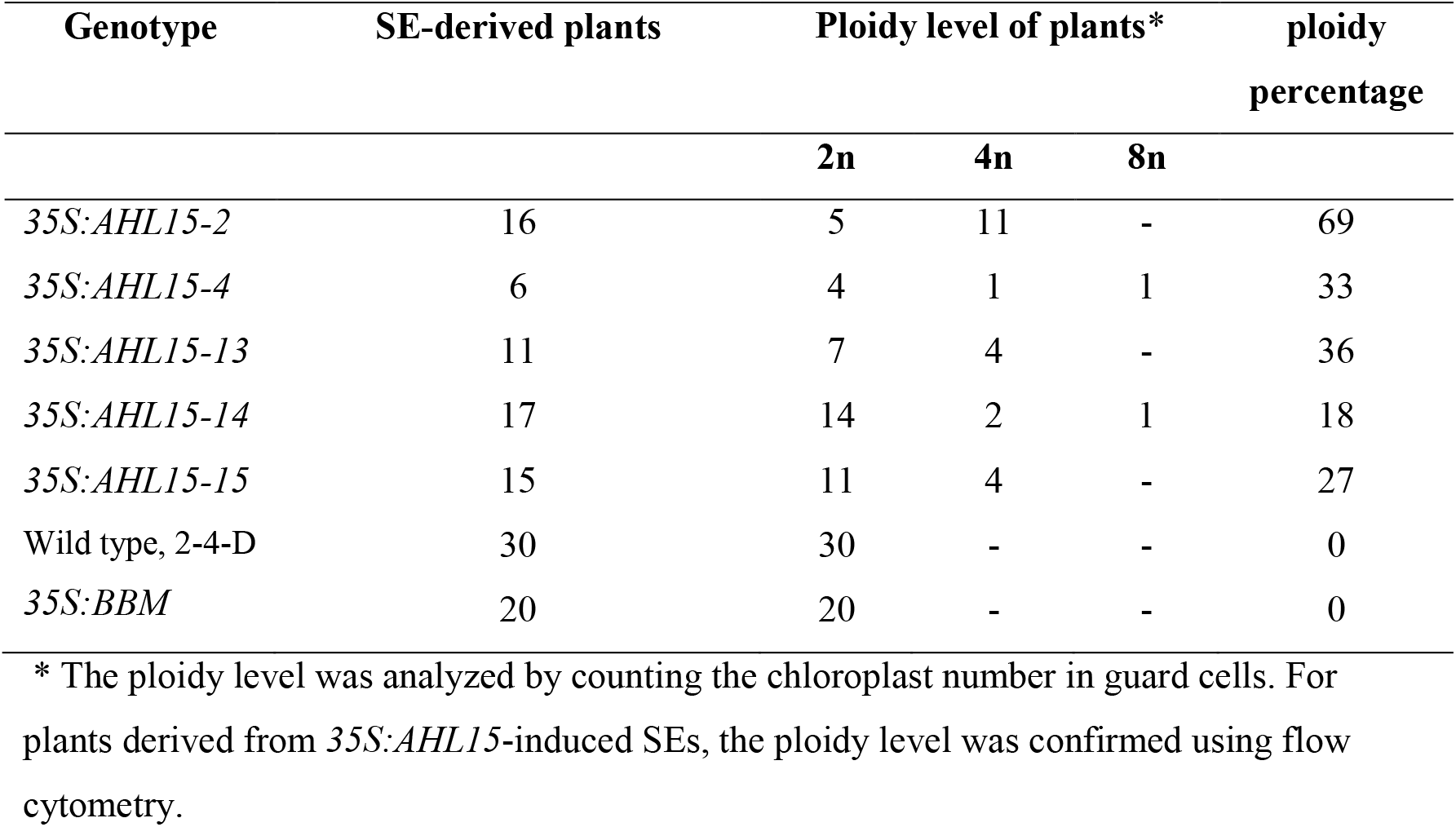
Ploidy level of plants derived from SEs induced by *AHL15* overexpression, *BBM* overexpression or by 2,4-D treatment

The considerable frequency of polyploid plants regenerated from *35S:AHL15* somatic embryos posed the question as to when polyploidisation is induced, and whether it is correlated with, or even promoted by SE induction. We observed a variable number of CENH3-GFP labelled centromeric dots (6-8, 12-15 and 25-30) per cell in cotyledons of *35S:AHL15* IZEs seven to eight days after of the start of culture, coinciding with SE induction and reflecting the presence of diploid, tetraploid and octoploid cells (Figure 5a). No evidence was obtained for polyploidy in root meristems (Figure S8a) or young leaves (Figure S8b) of *35S:AHL15* plants derived from ZEs, nor was polyploidy observed in the 2,4-D-induced non-embryogenic calli found on leaf and root tissues of *35S:AHL15* plants (Figure S8c, d). When we followed the *pH2B:H2B-GFP* reporter in cotyledons of *35S:AHL15* IZEs in time, we did not observe any cells with an increased number of chromocenters during the first week of culture (Figure 5b). At eight days of IZE culture, however, an increase in chromocenter number could be detected in proliferating *35S:AHL15* cotyledon cells (Figure 5c). This result showed that polyploidisation in *35S:AHL15* cotyledon cells is tightly associated with the induction of SE. Based on these results, and in line with the observation that *35S:AHL15* polyploid plants were only obtained from *35S:AHL15* somatic embryos, we conclude that the *AHL15*-induced polyploidisation occurs specifically during somatic embryo initiation.

**Figure 5.**
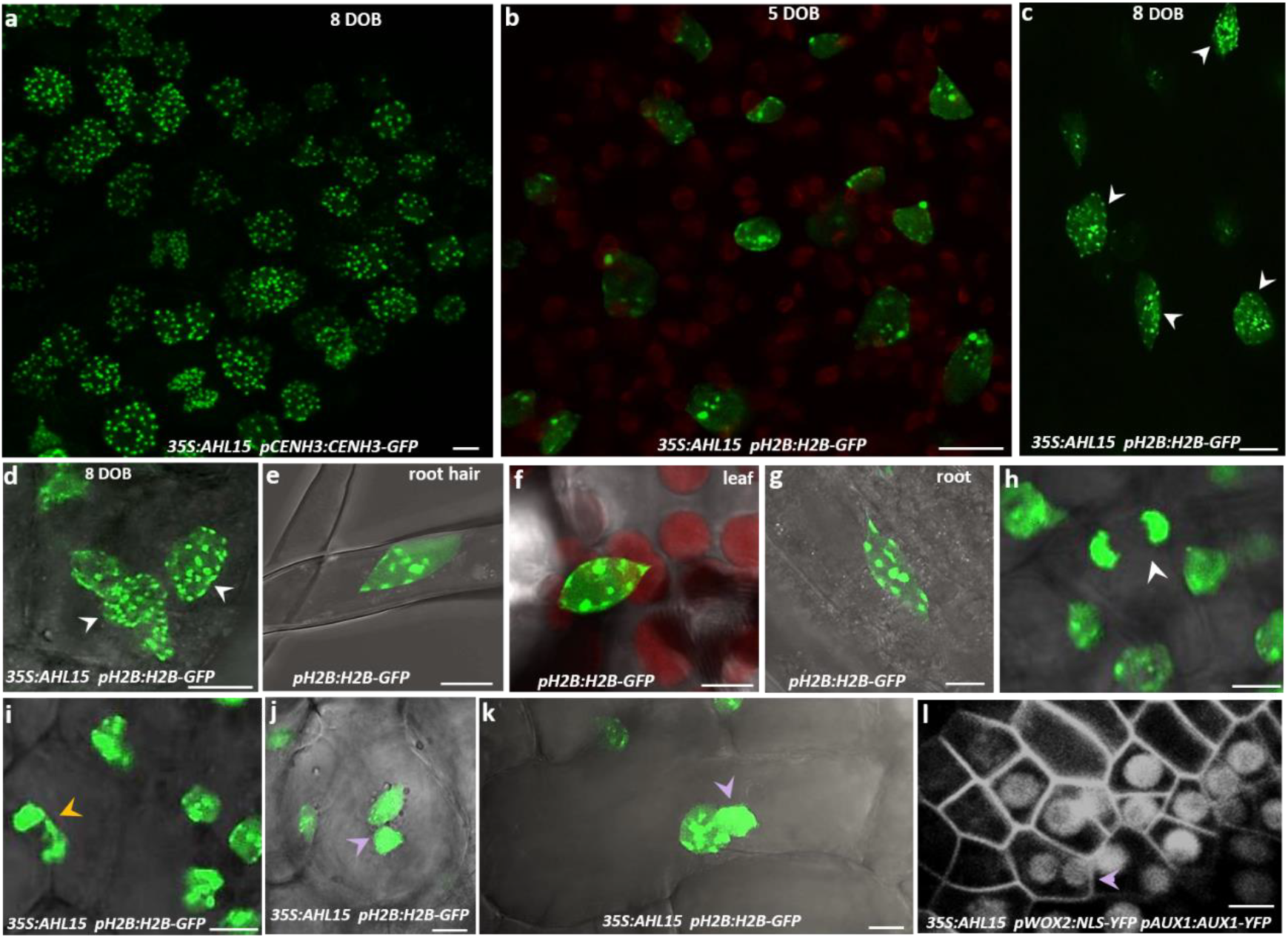
*AHL15* overexpression-induced SE results in polyploidy by endomitosis due to chromosome mis-segregation. (a) Confocal image of polyploid cells detected by CENH3-GFP-mediated centromere labeling in an embryonic structure developing on a cotyledon of a *35S:AHL15* IZE cultured for 8 days on B5 medium (DOB). (b,c) H2B-GFP-labelled chromocenters in nuclei of cotyledon cells of *35S:AHL15* IZEs cultured for 5 (b) or 8 (c) DOB. (d-g) Confocal images of H2B-GFP labelled chromocenters in nuclei of cotyledon cells of a *35S:AHL15* IZE cultured for 8 DOB (d) or in endoreduplicated nuclei of wild-type root hair (e), leaf (f), or root epidermis (g) cells. White arrowheads in (c) and (d) indicate cells with a duplicated number of chromocenters. (h-k) Confocal microscopy analysis of chromosome segregation dividing epidermis cells in cotyledons of *35S:AHL15* IZEs using the *pH2B:H2B-GFP* reporter. The white arrowhead indicates normal chromosome segregation during anaphase (h), the yellow arrowhead indicates mis-segregation of chromosomes during anaphase (i), and the magenta arrowhead indicates a bi-nucleated cell (j,k) in cotyledons of *35S:AHL15* IZEs cultured for 8 DOB. (l) Confocal microscopy image of a cotyledon of a *35S:AHL15 pWOX2:NLS-YFP pAUX1:AUX1-YFP* IZE. *pWOX2:NLS-YFP* and *pAUX1:AUX1-YFP* reporters were used to mark embryonic nuclei and plasma membranes, respectively. The magenta arrowhead indicates a bi-nucleated cell in an area of cells with WOX2-YFP-marked embryo cell fate. Images show a merge of the transmitted light and the GFP channel (d-k), or the GFP (a-c) or YFP (l) channel alone. Size bars indicate 6 µm.

### *AHL15* overexpression-induced polyploidy occurs by endomitosis due to chromosome mis-segregation

Endoreduplication normally occurs in expanding cells to facilitate cell growth. During endoreduplication duplicated chromosomes do not enter into mitosis and the number of chromocenters does not increase ^39–41^. We observed an increase in H2B-GFP-marked chromocenters in *35S:AHL15* cotyledon cells that coincided with polyploidisation events (Figure 5c and d), but not in endoreduplicated nuclei of wild-type root hair, leaf or root cells (Figure 5e-g), suggesting that these polyploid *35S:AHL15* cells are not derived from endoreduplication. Thus duplication of segregated chromosomes in *35S:AHL15* cotyledons cells must be caused by endomitotis, during which mitosis is initiated and chromosomes are separated, but cytokinesis fails to occur. Although ectopic overexpression of *AHL15* resulted in a high percentage of polyploid plants (Table 1), polyploidy of the embryo itself was not a prerequisite for further development of somatic embryos into plants, as most of the *AHL15* overexpressing plants were still diploid.

Disruption of heterochromatin in human mitotic cells leads to mis-segregation of chromosomes ^42–45^ and cellular polyploidization ^46^. We hypothesized that heterochromatin disruption and more global chromatin decondensation in dividing *35S:AHL15* cotyledon cells might contribute to endomitosis resulting in polyploid somatic embryo progenitor cells. Compared to chromosome segregation in wild-type dividing cotyledon cells (Figure 5h), chromosome segregation in dividing *35S:AHL15* cotyledon cells lagged behind (Figure 5i) and binucleate cells (Figure 5j-l) could be detected in cotyledons of 7-day-old *35S:AHL15* explants. The observation that binucleate cotyledon cells expressed the *pWOX2:NLS-YFP* embryo marker (Figure 5l) confirmed that such cells can adopt embryo identity and develop into polyploid somatic embryos. Taken together, we conclude that heterochromatin disruption in *35S:AHL15*-induced embryonic cotyledon cells leads to chromosome mis-segregation, the formation of binucleate cells and finally to cellular polyploidization coinciding with the development of polyploid somatic embryos.

## Discussion

The herbicide 2,4-D is extensively used for SE induction in a wide range of plant species, including *Arabidopsis*. In *Arabidopsis*, SE can also be induced on IZEs or seedlings in the absence of 2,4-D treatment by the overexpression of specific transcription factors, such as the AIL transcription factor BBM ^13,29^. In this study, we showed that AHL15 adds to the list of nuclear proteins whose overexpression induces somatic embryos on IZEs and seedlings in the absence of 2,4-D. In line with this observation, *AHL15* and several of its homologs are upregulated and required for SE induction upon 2,4-D treatment. Furthermore, they are required for efficient *BBM*-induced SE as downstream targets of BBM.

AT-hook motif-containing proteins are generally considered to be chromatin architecture factors ^17,30–32,47^. Studies in animals have shown that chromatin decondensation precedes the induction of pluripotent stem cells and their subsequent differentiation^48^. In the *Arabidopsis* zygote, predominant decondensation of the heterochromatin configuration is likely to contribute to the totipotency of this cell ^49^. Our data indicate that *AHL15* overexpression induces a global reduction of the amount of heterochromatin in induced somatic embryonic cells, whereas *ahl* loss-of-function mutants show enhanced heterochromatin formation in *in vitro* cultured explants, correlating with a reduced embryonic competence of their explants. Based on our results, we suggest a model in which chromatin opening is required for the acquisition of embryonic competence in somatic plant cells (Figure 6). In this model chromatin opening is mediated by upregulation of *AHL* gene expression, which can be achieved by *35S* promotor-driven overexpression, by 2,4-D treatment or by *BBM* overexpression.

**Figure 6.**
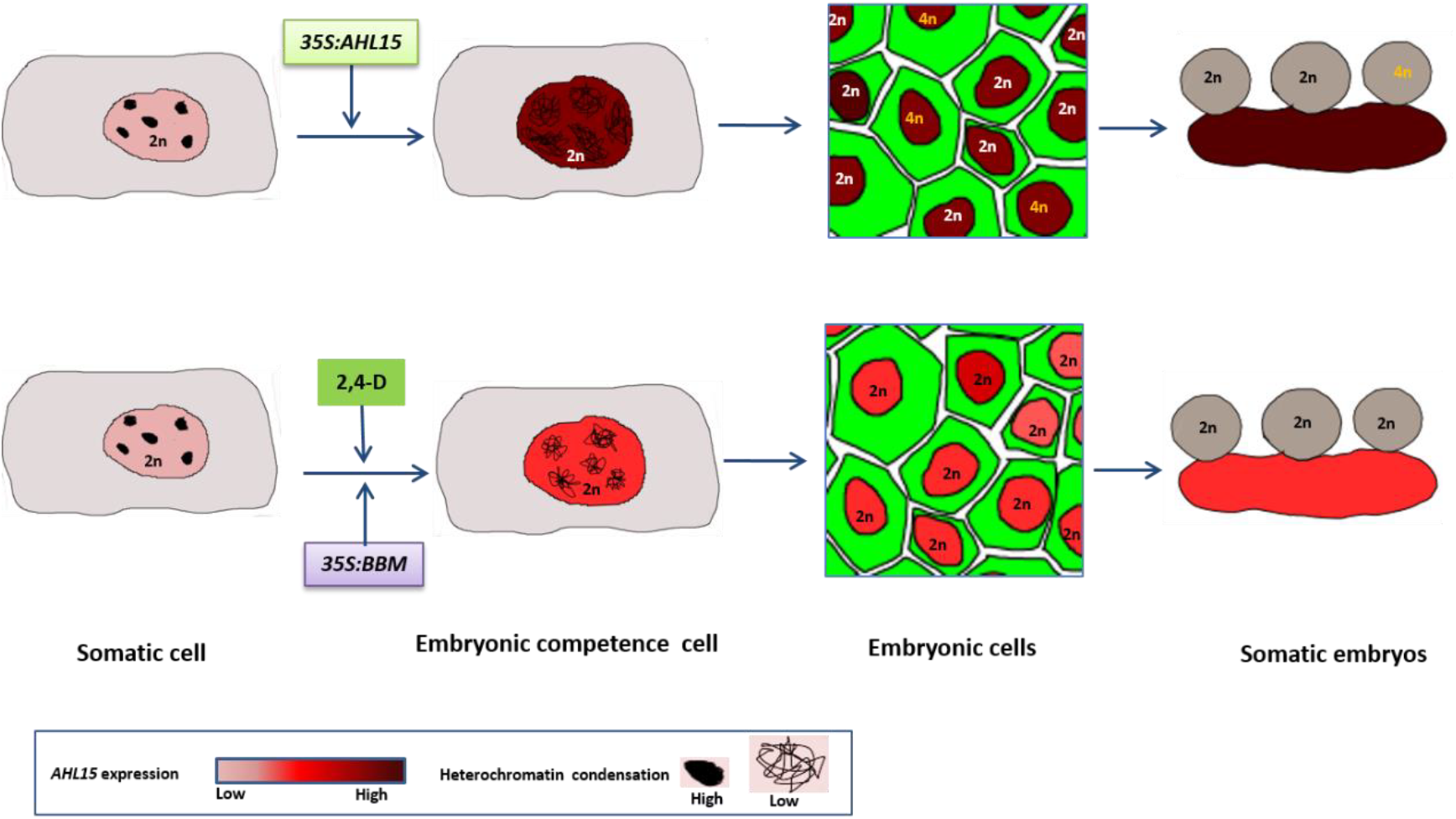
Model for somatic-to-embryonic reprogramming and cellular polyploidization by *AHL15* overexpression. *AHL15* or *BBM* overexpression, or 2,4-D treatment all induce the chromatin decondensation that is required to induce embryonic competence in somatic cells. The high level of heterochromatin decondensation obtained by *35S* promoter-driven *AHL15* overexpression prevents chromosome segregation in some cells, leading to endomitosis events that give rise to polyploid embryonic cells and subsequently to polyploid somatic embryos. *BBM* overexpression or 2,4-D treatment also lead to enhanced *AHL15* expression resulting in chromatin decondensation sufficient to induce embryonic cells and the resulting somatic embryos, but insufficient to leading to endomitosis, thus only giving rise to diploid somatic embryos.

During the S phase of the cell cycle, eukaryotic cells duplicate their chromosomes after which the mitosis machinery ensures that the sister chromatids segregate equally over the two daughter cells. However, some cell types do not separate the duplicated chromosomes, leading to a polyploidy state known as endopolyploidy ^50^. In plants, endopolyploidy is commonly classified either as endomitosis or endoreduplication ^50^. Endoreduplication occurs during cellular differentiation, where chromosomes are duplicated but do not segregate, leading to the formation of polytene chromosomes ^40^. By contrast, during endomitosis sister chromatids are separated, but the last steps of mitosis including nuclear division and cytokinesis are skipped, generally leading to a duplication of the chromosome number. In this work we show that polyploid cells can be specifically detected during *35S:AHL15* induced SE.

Previous studies have shown that defects in heterochromatin condensation in animal cells lead to mis-separation of chromosomes during mitosis ^42–45^, and that mis-segregation of chromosomes subsequently leads to cellular polyploidisation ^46^. In our experiments, we found a high reduction of heterochromatin coinciding with mis-segregation of chromosomes in *in vitro*-cultured *35S:AHL15* cotyledon cells. Consistent with the strong conservation of chromosome segregation mechanisms between animal and plant cells ^51^, we propose that cellular polyploidisation in *35S:AHL15* embryonic cells is caused by an ectopic AHL15-mediated reduction in chromosome condensation during mitosis, which results in chromosome mis-segregation. The observation that polyploid embryos and plants are not obtained after 2,4-D treatment or by *BBM* overexpression suggests that in these somatic embryos *AHL15* expression levels are finely balanced to prevent chromosome mis-segregation (Figure 6).

A low frequency of polyploid plants derived from somatic embryo cultures has been reported ^8,52–55^, but the molecular and cytological basis for this genetic instability in relation to *in vitro* embryogenesis has not been described. Since SE requires cell division, these genome duplications most likely have been caused by endomitosis and not by endoreduplication. Until now, endomitosis in plants has been described as a result of defective cytokinesis due to aberrant spindle- or cell plate formation ^38,56,57^. Our results show that chromosome mis-segregation by *AHL15* overexpression-induced chromatin decondensation provides an alternative mechanism of endomitosis in plants, possibly acting not only during SE but also during gametogenesis or zygotic embryogenesis as infrequent environmentally-induced events that could lead to genome duplication-enabled speciation during evolution. Environmentally-induced AHL15-mediated genome duplication might also provide an alternative method for chromosome doubling in cultured embryos derived from haploid explants, such as egg cells or microspores ^58^ and for the production of polyploid crops.

## Methods

### Plant material and growth conditions

T-DNA insertion mutants *ahl15* (SALK_040729) and *ahl19* (SALK_070123) were obtained from the European Arabidopsis Stock Centre (http://arabidopsis.info/). Primers used for genotyping are described in Table S1. The reporter lines p*CENH3:CENH3-GFP*^37^ and p*H2B:H2B-GFP* ^37^, *pWOX2::NLS-YFP* ^59^, and *pAUX1:AUX1-YFP* ^60^ have been described previously. For *in vitro* plant culture, seeds were sterilized in 10 % (v/v) sodium hypochlorite for 12 minutes and then washed four times in sterile water. Sterilized seeds were plated on MA medium ^61^ containing 1 % (w/v) sucrose and 0.7 % agar. Seedlings, plants and explants were grown at 21°C, 70% relative humidity, and 16 hours photoperiod.

### Plasmid construction and plant transformation

The *35S:AHL15* construct was generated by PCR amplification of the full-length *AHL15* cDNA of (AT3G55560) from ecotype Columbia (Col-0) using primers 35S:AHL15-F and -R (Table S1). The resulting PCR product was cloned as a *Sma*I/*Bgl*II fragment into the *p35S-3’OCS* expression cassette of plasmid pART7, which was subsequently cloned as *Not*I fragment into the binary vector pART27 ^62^. To generate the other overexpression constructs, the full-length cDNA clones of *AHL19* (AT3G04570), *AHL20* (AT4G14465), and *AHL29* (AT1G76500) from *Arabidopsis* Col-0 were used to amplify the open reading frames (ORFs) using primers indicated in Table S2. The ORFs were cloned into plasmid pJET1/blunt (GeneJET™ PCR Cloning Kit, #K1221), and next transferred as *Not*I fragments to binary vector *pGPTV 35S:FLAG* ^63^. To generate the *pAHL15:AHL15-GUS* and *pAHL15:AHL15-TagRFP* translational fusions, a 4 kb fragment containing the promoter and the full coding region of *AHL15* was amplified using PCR primers AHL15-GUS-F and -R (Table S1), and inserted into pDONR207 using a BP reaction (Gateway, Invitrogen). LR reactions were carried out to fuse the 4 kb fragment upstream of GUS and *tagRFP* in respectively destination vectors pMDC163 ^64^ and pGD121 ^65^. The artificial microRNA (amiR) targeting *AHL20* was generated as described by Schwab and colleagues ^66^ using oligonucleotides I-IV miR-a/s AHL20 (Table S1). The fragment of the *amiRAHL20* precursor was amplified using PCR primers amiRNA AHL20-F and -R (Table S1), and subsequently introduced into the entry vector pDONR207 via a BP reaction (Gateway, Invitrogen). The *amiRAHL20* precursor was recombined into destination vectors pMDC32 ^64^ downstream of the *35S* promoter via an LR reaction (Gateway, Invitrogen). To generate the *pAHL15:AHL15-ΔG* construct, a synthetic *Kpn*I-*Spe*I fragment containing the *AHL15* coding region lacking the sequence encoding the Gly-Arg-Phe-Glu-Ile-Leu amino acids in the C-terminal region (BaseClear) was used to replace the corresponding coding region in the *pAHL15:AHL15* construct. The *p35S:BBM-GR* construct has been described previously ^67^. All binary vectors were introduced into *Agrobacterium tumefaciens* strain AGL1 by electroporation^68^ and transgenic *Arabidopsis* Col-0 lines were obtained by the floral dip method ^69^.

The *p35S:BBM-GR* construct has been described previously^67^. All binary vectors were introduced into *Agrobacterium tumefaciens* strain AGL1 by electroporation^68^ and transgenic *Arabidopsis* Col-0 lines were obtained by the floral dip method ^69^.

### Somatic embryogenesis

Immature zygotic embryos (IZEs) at the bent cotyledon stage of development (10-12 days after pollination) or germinating dry seeds were used as explants to induce SE using a previously described protocol ^8^. In short, seeds and IZEs were cultured on solid B5 medium^70^ supplemented with 5 μM 2,4-D, 2 % (w/v) sucrose and 0.7 % agar (Sigma) for 2 weeks. Control seeds or IZEs were cultured on solid B5 medium without 2,4-D. To allow further embryo development, explants were transferred to medium without 2,4-D. One week after subculture, the capacity to induce SE was scored under a stereomicroscope as the number of somatic embryos produced from 50 explants cultured on a plate. Three plates were scored for each line. The Student’s *t*-test was used for statistical analysis of the data.

### Quantitative real-time PCR (RT-qPCR) and ChIP seq analysis

To determine the expression of *AHL* genes during SE induction, RNA was isolated from 25 IZEs cultured for 7 days on B5 medium with or without 2,4-D in 4 biological replicates using a Qiagen RNeasy Plant Mini Kit. The RNA samples were treated with Ambion^®^ TURBO DNA-free™ DNase. To determine the expression of *AHL* genes in 2,4-D treated Col-0 IZEs by qRT-PCR, 1 µg of total RNA was used for cDNA synthesis with the iScript™ cDNA Synthesis Kit (BioRad). PCR was performed using the SYBR-Green PCR Master mix (Biorad) and a CHOROMO 4 Peltier Thermal Cycler (MJ RESEARCH). The relative expression level of *AHL* genes was calculated according to the 2^−ΔΔCt^ method ^71^, using the without 2,4-D value to normalize and *β-TUBULIN-*6 (At5g12250) gene as a reference. The gene-specific PCR primers used are described in Table S1.

The effect of *BBM* overexpression on *AHL* gene expression was examined by inducing five-day-old *Arabidopsis thaliana* Col-0 and *35S:BBM-GR* seedlings (four biological replicates for each line) for three hours with 10 µM dexamethasone (DEX) plus 10 µM cycloheximide (CHX). RNA was isolated using the Invitek kit, treated with DNAseI (Invitrogen) and then used for cDNA synthesis with the Taqman cDNA synthesis kit (Applied Biosystems). qRT-PCR was performed as described above. The relative expression level of *AHL* genes was calculated according to the 2^−ΔΔCt^ method ^71^, using the wild-type Col-0 value to normalize and the *SAND* gene (At2g28390; ^72^ as a reference. The gene-specific PCR primers are listed in Table S1.

The ChIP-seq data and analysis was downloaded from GEO (GSE52400). Briefly, the experiments were performed using somatic embryos from either 2,4-D-induced *BBM:BBM-YFP* cultures (with *BBM:NLS-GFP* as a control) or a *35S:BBM-GFP* overexpression line (with *35S:BBM* as a control), as described in ^73^.

### Ploidy analysis

The ploidy level of plants derived from *35S:AHL15*-induced somatic embryos was determined by flow cytometry (Plant Cytometry Services, Schijndel, Netherlands), and confirmed by counting the total number of chloroplasts in stomatal guard cells and by comparing flower size and or the size of the nucleus in root epidermal cells. The number of chloroplasts in stomatal guard cells was counted for plants derived from 2,4-D- and BBM-induced somatic embryos.

### Histological staining and microscopy

Histochemical β-glucuronidase (GUS) staining of *pAHL15:AHL15-GUS* IZEs or ovules was performed as described previously^74^ for 4 hours at 37 °C, followed by rehydration in a graded ethanol series (75, 50, and 25 %) for 10 minutes each. GUS stained tissues were observed and photographed using a LEICA MZ12 microscopy (Switzerland) equipped with a LEICA DC500 camera.

DNA staining of wild-type and *35S:AHL15* seedlings was performed using propidium iodide (PI) according to the protocol described by Baroux *et al*. ^75^. To stain nuclei, the samples were incubated for 30 minutes in 4′,6-diamidino-2-phenylindole (DAPI) staining solution (1 μg/ml DAPI in phosphate-buffered saline (PBS) just before observation.

For scanning electron microscopy (SEM), seedlings were fixed in 0.1 M sodium cacodylate buffer (pH 7.2) containing 2.5% glutaraldehyde and 2% formaldehyde. After fixation, samples were dehydrated by a successive ethanol series (25, 50, 70, 95, and 100 %), and subsequently critical-point dried in liquid CO_2_. Dried specimens were gold-coated and examined using a JEOL SEM-6400 (Japan).

For morphological studies of embryos, fertilized ovules were mounted in a clearing solution (glycerol:water:chloral hydrate = 1:3:8 v/v) and then incubated at 65 °C for 30 min and observed using a LEICA DC500 microscopy (Switzerland) equipped with differential interference contrast (DIC) optics.

The number of chloroplasts in leaf guard cells, the size of the DAPI stained nuclear area in root cells and the number of conspicuous heterochromatin regions of the PI stained nuclei of cotyledon cells were recorded using a confocal laser scanning microscope (ZEISS-003-18533), using a 633 laser, a 488 nm LP excitation and a 650-700 nm BP emission filters for chlorophyll signals in guard cells, a 405 laser, a 350 nm LP excitation and a 425-475 nm BP emission filters for DAPI signals in cotyledon cells, and a 633 laser, 488 nm LP excitation and 600-670 nm emission BP filters for PI signals in cotyledon cells. The relative size chromocenter spots were measured from confocal images by measuring of region of the spots using the measuring region tool of ImageJ software (Rasband).

Cellular and subcellular localization of AHL15-TagRFP and H2B- or CENH3-GFP protein fusions were visualized using the same laser scanning microscope with a 633 laser, and a 532 nm LP excitation and 580-600 nm BP emission filters for TagRFP signals and a 534 laser, 488 nm LP excitation and 500-525 nm BP emission filters for GFP signals.

## Author contribution

B.vd.Z. and R.O. conceived the project, O.K., M.C., A.H., K.B., B.vd.Z. and R.O. designed experiments, and analyzed and interpreted results, O.K. performed the majority of the experiments, A.R. contributed to polyploidisation analysis and generated *pAHL15:AHL15-ΔG* transgenic lines, P.M. generated and analysed the *Arabidopsis* lines overexpressing the AHL genes, A.H. performed the BBM-related experiments, O.K. and R.O. wrote the manuscript, K.B., B.vd.Z. and A.H. assisted in finalizing the manuscript.

## Acknowledgements

We thank Gerda Lamers for help with microscopy and are grateful to Nick Surtel, Ward de Winter and Jan Vink for their help with plant growth and media preparation. O.K. was supported in part by Grant G14_006.02 to R.O. from Generade, and by Building Blocks of Life grant 737.016.013 to R.O. from the Netherlands Organisation for Scientific Research. M.C. and P.M. we partially supported through grant IS054064 from the Ministry of Economic Affairs (SenterNovem). The authors declare no competing financial interests.

