## Supplemental figures for "An *Arabidopsis* AT-hook motif nuclear protein mediates somatic embryogenesis and coinciding genome duplication"

**Supplemental Information**

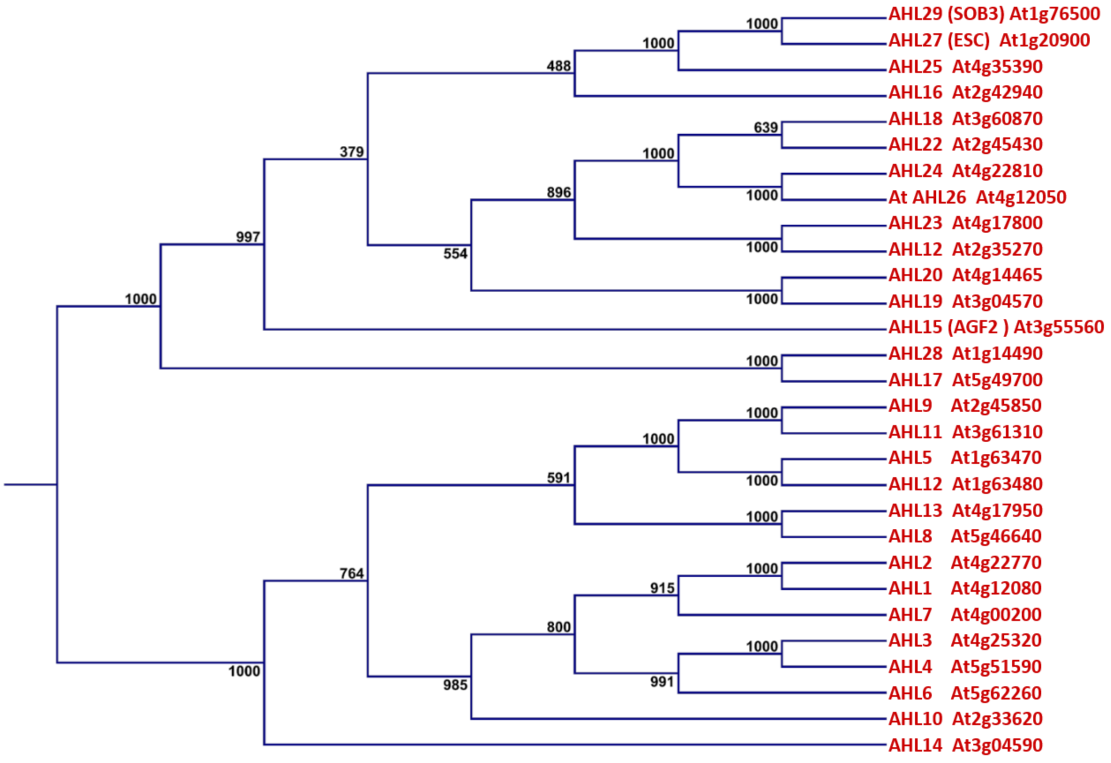


**Figure S1. A phylogenetic tree of the *Arabidopsis AHL* gene family.**

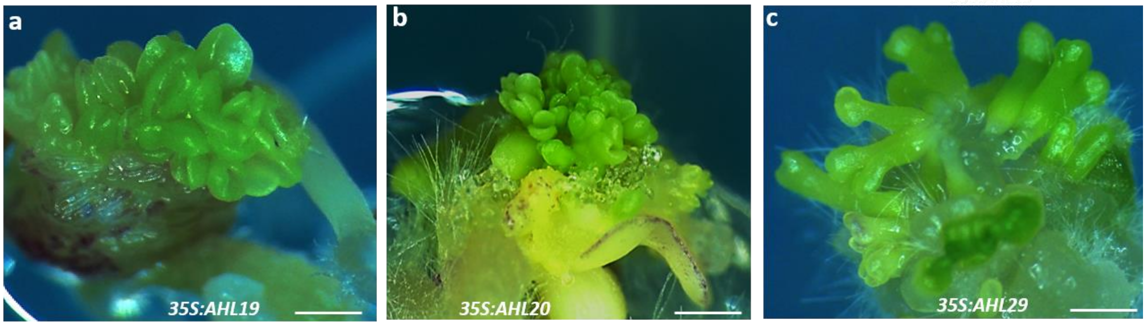


**Figure S2. Overexpression of *AHL19, AHL20 and AHL20* induces SE.** (a-c) The embryo structures induced on IZEs of *35S:AHL19* (a), *35S:AHL20* (b) or *35S:AHL29* (c) plants cultured for 2 weeks on medium lacking 2,4-D. Size bar indicates 1 mm.

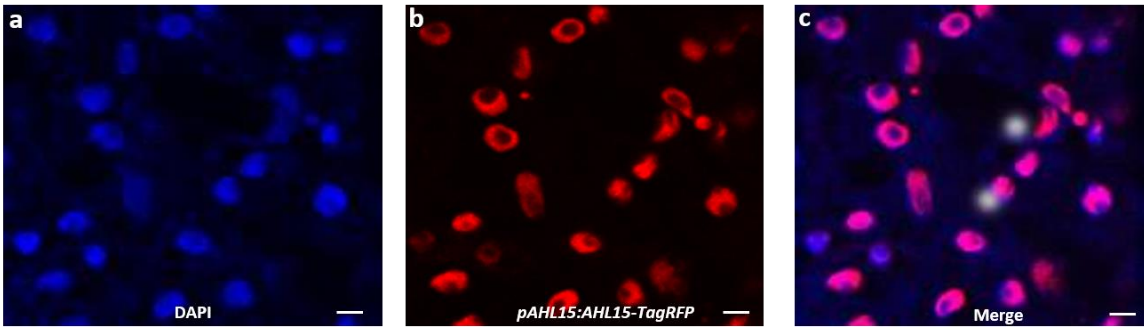


**Figure S3. Nuclear localization of AHL15 in embryo cells.** (a-c) Confocal images of embryo cells in torpedo stage. The blue channel showing nuclear staining by DAPI (a), the RFP channel showing nuclear-localized AHL15-tagRFP (b), and the merged images (c). Size bar indicates 4 µm.

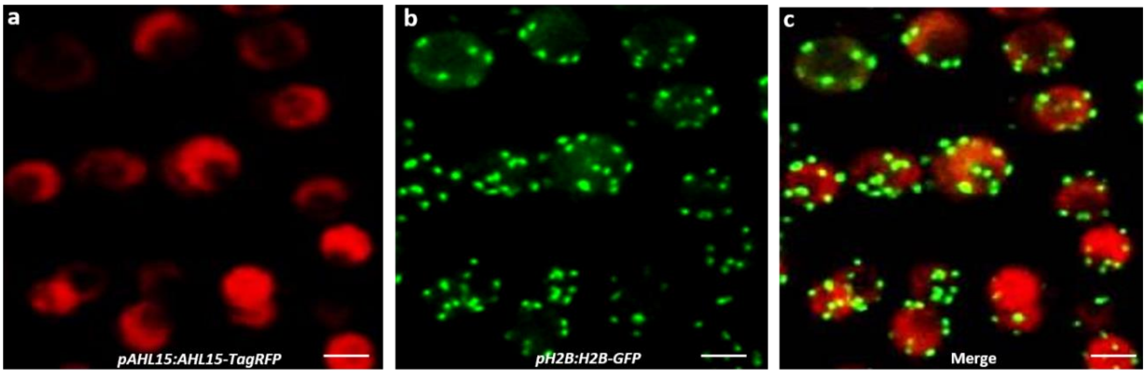


**Figure S4. AHL15-TagRFP does not colocalize with H2B-GFP-marked heterochromatin.** (a-c) Confocal images of root meristem cells. The RFP channel showing nuclear-localised AHL15-tagRFP (a), the GFP channel showing H2B-GFP marked heterochromatin (b), and the merged image (c). Size bar indicates 5 µm.

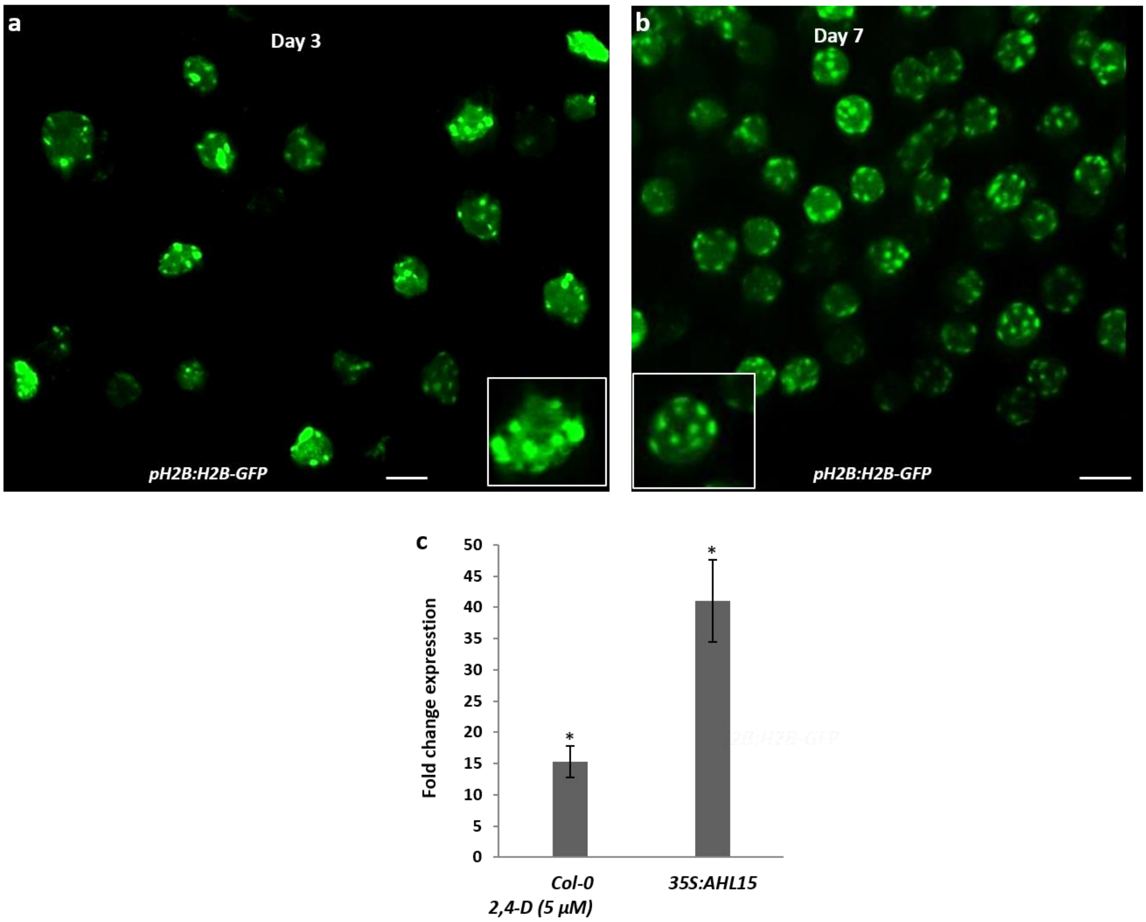


**Figure S5. 2,4-D treatment does not lead to substantial heterochromatin decondensation.** Condensed heterochromatin visualized by H2B-GFP labelled nuclei in cotyledon cells of *H2B:H2B-GFP* IZEs, 3 (a) or 7 (b) days after culture on B5 medium containing 2,4-D. Size bar indicates 5 µm. (c) Enhanced *AHL15* expression in both wild-type 2,4-D treated IZEs and in *35S:AHL15* IZEs for 7 days compared to wild-type IZEs on B5 medium without 2,4-D. Error bars indicate the standard error of the mean of three biological replicates. Asterisks indicate a significant enhancement of *AHL15* gene expression in *35S:AHL15* IZEs cultured without 2,4-D  or in IZEs cultured on medium with 2,4-D and compared to medium without 2,4-D (Student's *t*-test, $p < 0.001$).

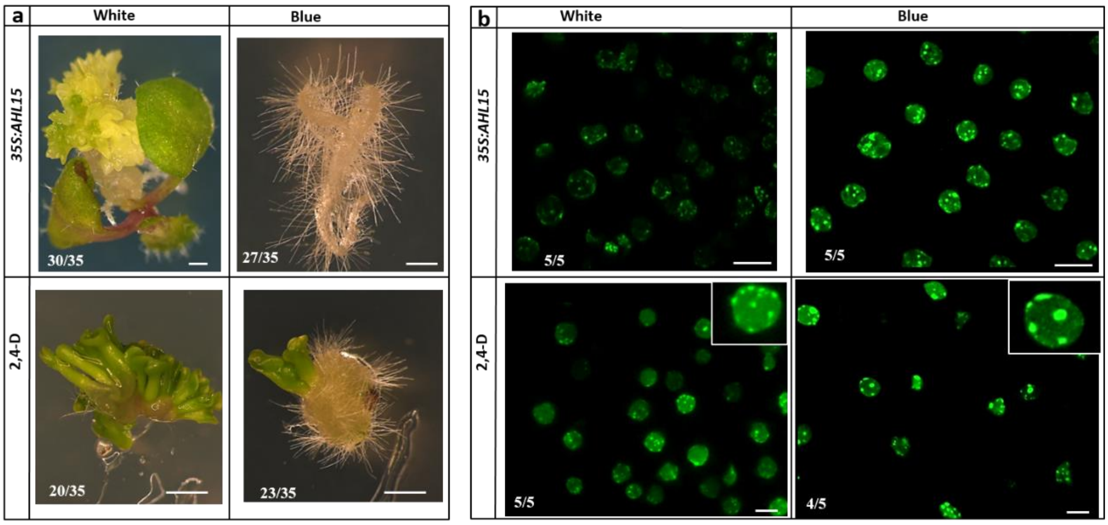


**Figure S6 | Blue light treatment inhibits SE and heterochromatin decondensation.** (a) The morphology of somatic embryo structures on *35S:AHL15* IZEs cultured on B5 medium (upper panel) or on wild-type IZEs cultured on B5 medium containing 2,4-D (lower panel) under white (left) or blue (right) light conditions. (b) H2B-GFP labelled nuclei in cotyledon cells of *35S:AHL15* or wild-type IZEs 7 days after culture on B5 medium (top panels) or on B5 medium containing 2,4-D (low panels), respectively, under white (left) or blue (right) light conditions. The plates containing IZEs were grown under blue (450 nm, 120 µmol $m^{-2}$ $s^{-1}$), or white (120 µmol $m^{-2}$ $s^{-1}$) LED lights. Size bar indicates 1 mm in a and 4 µm in b.

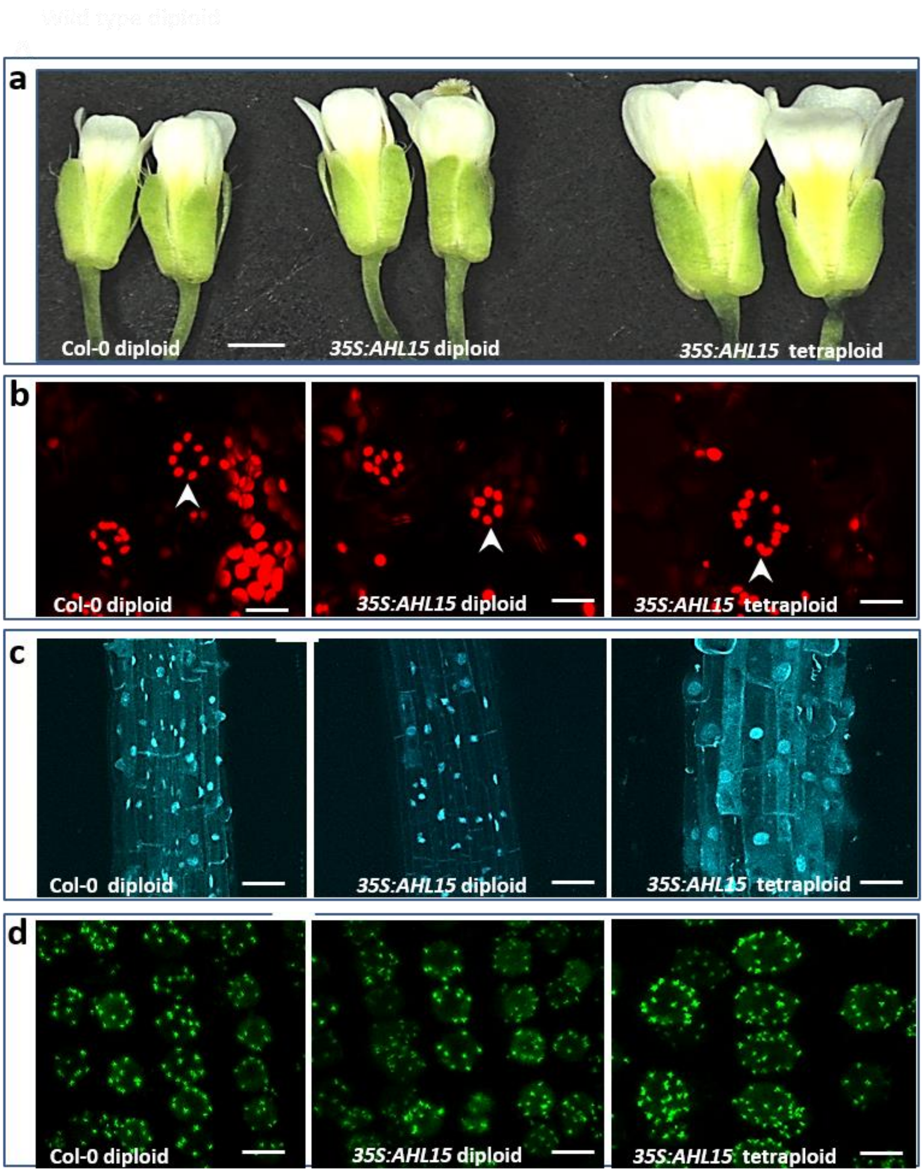


**Figure S7. Plants regenerated from *35S:AHL15*-induced somatic embryos are frequently polyploid.** (a-d) Analysis of wild-type *Arabidopsis* (left), and a diploid plant line (middle), and a tetraploid plant line (right) each regenerated from an *35S:AHL15*-induced somatic embryo. (a) Tetraploid *35S:AHL15* plants show increased organ size compared to the diploid control plants, as demonstrated by the size of the flower organs. (b-d) Tetraploid *35S:AHL15* plants have twice the number/a higher number of chloroplasts in guard cells (marked by arrow heads, b), show root cells with a larger nucleus and cell size (c), and show a duplication in the CENH3-GFP-labelled centromeres (d) compared to wild-type and diploid *35S:AHL15 Arabidopsis* plants. Size bar indicates 1mm in A, 8 µm in b, 22 µm in c and 6 µm in d.

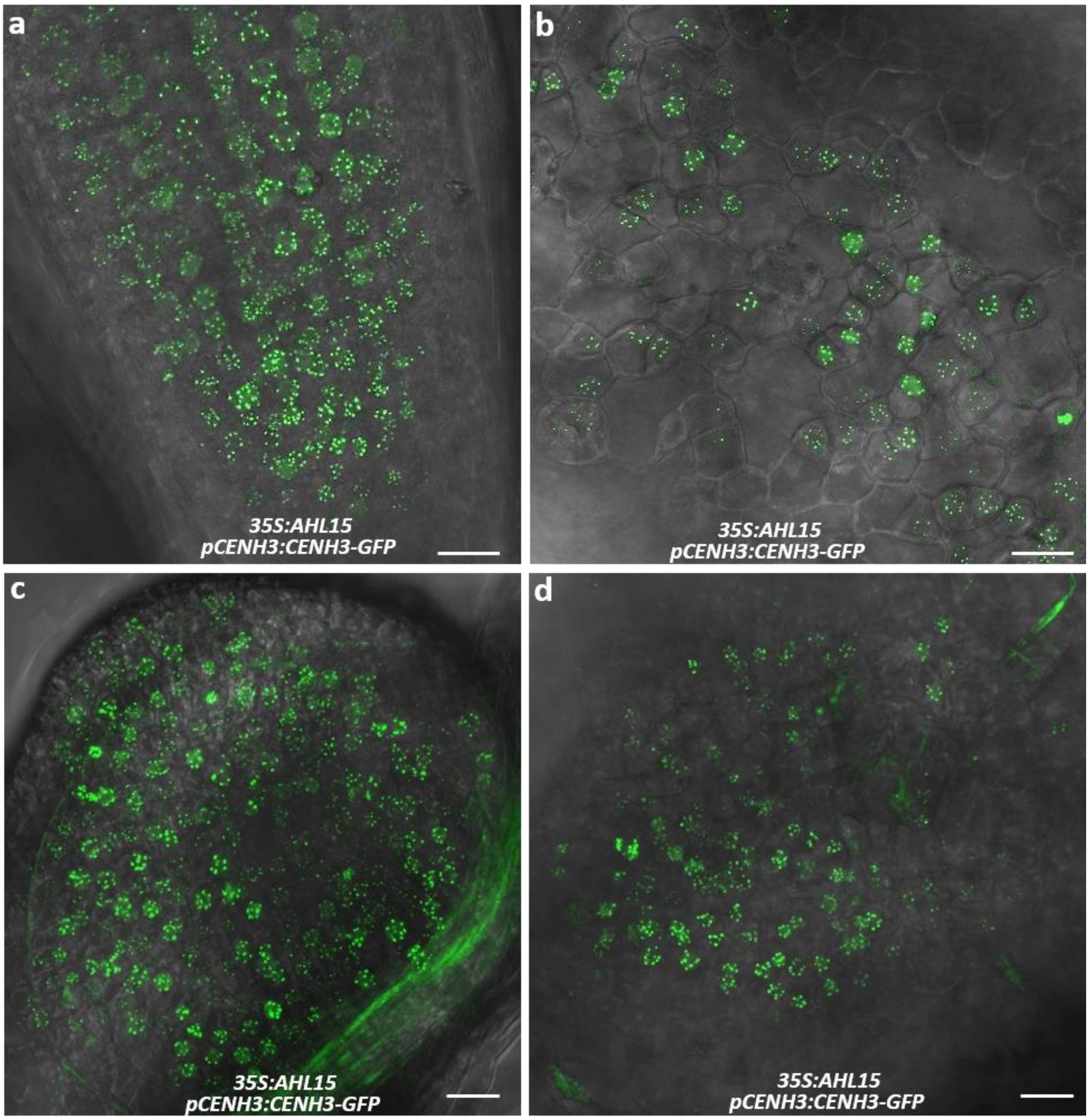


**Figure S8. Endoploidy does not occur in *35S:AHL15* leaves and roots.** (a-d) CENH3-GFP–mediated centromere labeling in cells of a root tip (a), a young leaf (b), or in cells of 2,4-D-induced callus on a root (c), or a young leaf (d). Size bar indicates 12 µm.

**Table S1.** Primers used for cloning, genotyping and qRT-PCR (F: forward; R: reverse)

| Name | Sequence (5' to 3') | Purpose |
|---|---|---|
| 35S:AHL15-F | CCCGGGATGGCGAATCCTTGGTGGGTAG | *35S:AHL15 construct* |
| 35S:AHL15-R | GGATCCTCAATACGAAGGAGGAGCACG | |
| 35S:AHL29-F | ATAAGAATGCGGCCGCGACGGTGGTTACGATCAATC | *35S:AHL29* construct |
| 35S:AHL29-R | ATAGTTTAGCGGCCGCCTAAAAGGCTGGTCTTGGTG | |
| 35S:AHL20 –F | ATAAGAATGCGGCCGCGCAAACCCTTGGTGGACGAAC | *35S:AHL20* construct |
| 35S:AHL20-R | ATAGTTTAGCGGCCGCTCAGTAAGGTGGTCTTGCGT | |
| 35S:AHL19-F | GGGGACAAGTTTGTACAAAAAAGCAGGCTCGATGGCGAATCCATGGTGGAC | *35S:AHL19 construct* |
| 35S:AHL19-R | GGGGACCACTTTGTACAAGAAAGCTGGGTAAACAAGTAGCAACTGACTGG | |
| pAHL15-GUS-F | GGGGACAAGTTTGTACAAAAAAGCAGGCTCGACACTCCTCTGTGCCACATT | *pAHL15:AHL15-GUS* construct |
| pAHL15-GUS-R | GGGGACCACTTTGTACAAGAAAGCTGGGTAATACGAAGGAGGAGCACGAG | *pAHL15:AHL15-tagRFP* |
| I miR-s AHL20 | GATTAGACTACCTCAAATTGCTATCTCTCTTTTGTATTCC | |
| II miR-a AHL20 | GATAGCAATTTGAGGTAGTCTAATCAAAGAGAATCAATGA | |
| III miR*s AHL20 | GATAACAATTTGAGGAAGTCTATTCACAGGTCGTGATATG | *35S:amiRAHL20* construct |
| IV miR*a AHL20 | GAATAGACTTCCTCAAATTGTTATCTACATATATATTCCT | |
| amiRNA AHL20-F | GGGGACAAGTTTGTACAAAAAAGCAGGCTCGCGACGGTATCGATAAGCTTG | |
| amiRNA AHL20-R | GGGGACCACTTTGTACAAGAAAGCTGGGTACCCATGGCGATGCCTTAAAT | |
| SALK_040729-F | GTCGGAGAGCCATCAACACCA | ahl15 genotyping |
| SALK_040729-R | CGACGACCCGTAGACCCGGATC | |
| SALK_070123-F | GGCGAATCCATGGTGGACAGG | ahl19 genotyping |
| SALK_070123-R | GGCCGCTCATCTGTCCTCCTC | |
| qAHL15-F | AAGAGCAGCCGCTTCAACTA | qRT-PCR *AHL15* |
| qAHL15-R | TGTTGAGCCATTTGATGACC | |
| qAHL20-F | CAAGGCAGGTTTGAAATCTTATCT | qRT-PCR *AHL120* |
| qAHL20-R | TAGCGTTAGAGAAAGTAGCAGCAA | |
| qAHL19-F | CTCTAACGCGACTTACGAGAGATT | qRT-PCR *AHL19* |
| qAHL19-R | ATATTATACACCGGAAGTCCTTGGT | |
| qβ-TUBULIN-6-F | TGGGAACTCTGCTCATATCT | qRT-PCR *TUBULIN-6* |
| qβ-TUBULIN-6-R | GAAAGGAATGAG GTTCACTG | |
| qSAND-F | AACTCTATGCAGCATTTGATCCACT | qRT-PCR *SAND* |
| qSAND-R | TGATTGCATATCTTTATCGCCATC | |
